# Selective deletion of *Methyl CpG binding protein 2* from parvalbumin interneurons in the auditory cortex delays the onset of maternal retrieval in mice

**DOI:** 10.1101/2023.01.30.526321

**Authors:** Deborah D. Rupert, Alexa Pagliaro, Jane Choe, Stephen D. Shea

## Abstract

Mutations in *MECP2* cause the neurodevelopmental disorder Rett syndrome. *MECP2* codes for methyl CpG binding protein 2 (MECP2), a transcriptional regulator that activates genetic programs for experience-dependent plasticity. Many neural and behavioral symptoms of Rett syndrome may result from dysregulated timing and threshold for plasticity. As a model of adult plasticity, we examine changes to auditory cortex inhibitory circuits in female mice when they are first exposed to pups; this plasticity facilitates behavioral responses to pups emitting distress calls. Brain-wide deletion of *Mecp2* alters expression of markers associated with GABAergic parvalbumin interneurons (PVin) and impairs the emergence of pup retrieval. We hypothesized that loss of *Mecp2* in PVin disproportionately contributes to the phenotype. Here we find that deletion of *Mecp2* from PVin delayed the onset of maternal retrieval behavior and recapitulated the major molecular and neurophysiological features of brain-wide deletion of *Mecp2*. We observed that when PVin-selective mutants were exposed to pups, auditory cortical expression of PVin markers increased relative to that in wild type littermates. PVin-specific mutants also failed to show the inhibitory auditory cortex plasticity seen in wild type mice upon exposure to pups and their vocalizations. Finally, using an intersectional viral genetic strategy, we demonstrate that post-developmental loss of *Mecp2* in PVin of the auditory cortex is sufficient to delay onset of maternal retrieval. Our results support a model in which PVin play a central role in adult cortical plasticity and may be particularly impaired by loss of *Mecp2*.

**SIGNIFICANCE STATEMENT:** Rett syndrome is a neurodevelopmental disorder that includes deficits in both communication and the ability to update brain connections and activity during learning (‘plasticity’). This condition is caused by mutations in the gene *MECP2*. We use a maternal behavioral test in mice requiring both vocal perception and neural plasticity to probe *Mecp2’*s role in social and sensory learning. *Mecp2* is normally active in all brain cells, but here we remove it from a specific population (‘parvalbumin neurons’). We find that this is sufficient to delay learned behavioral responses to pups and recreates many deficits seen in whole brain *Mecp2* deletion. Our findings suggest that parvalbumin neurons specifically are central to the consequences of loss of *Mecp2* activity and yield clues as to possible mechanisms by which Rett syndrome impairs brain function.

## INTRODUCTION

Rett Syndrome (RTT) is a pervasive neurodevelopmental disorder that results from sporadic, *de novo* loss-of-function mutations in the *MECP2* gene, which codes for the transcriptional regulator methyl CpG binding protein-2 (Amir et al., 1999; Samaco et al., 2008). Because *MECP2* is located on the X chromosome, when it possesses a disabling mutation, males (or other individuals with a single X chromosome) lose their sole functioning copy and typically die perinatally; females (or other individuals with two X chromosomes) have heterozygous mosaic expression, and frequently survive infancy with impairments in cognition, musculoskeletal structure, metabolism (Van den Veyver and Zoghbi, 2000; Braunschweig et al., 2004), auditory processing, language, and communication (Bashina et al., 2002; Glaze, 2005). *MECP2* has been repeatedly implicated in the regulation of neural plasticity (Deng et al., 2010; McGraw et al., 2011; Noutel et al., 2011; Na et al., 2013; Deng et al., 2014; He et al., 2014; Krishnan et al., 2015; Tai et al., 2016; Krishnan et al., 2017; Gulmez Karaca et al., 2018). This observation, combined with the non-linear developmental course of RTT, has fueled speculation that *MECP2* is most essential during periods of elevated neuronal plasticity, e.g. early critical periods (He et al., 2014; Krishnan et al., 2015).

Mouse models in which *Mecp2* expression has either been disabled (Chen et al., 2001; Guy et al., 2001) or deleted with spatiotemporal selectivity using flanking loxP sites and cell type-specific expression of Cre recombinase (Gemelli et al., 2006) have been invaluable for understanding the biology of Mecp2. We capitalize on these models with a natural behavior, pup retrieval by female mice, as a readout of cortical plasticity and function (Krishnan et al., 2017; Lau et al., 2020). Our past work was performed in female mice that lacked one functional copy of *Mecp2* (*Mecp2^het^*). The mosaicism in these mice more closely represent the genetic condition in humans, as compared to the more commonly used male null model.

When female mice are first exposed to pups, they exhibit ‘pup retrieval’ (Rosenblatt, 1967; Sewell, 1970; Ehret et al., 1987), a learned behavioral response to the ultrasonic cries emitted by distressed or wandering pups (Galindo-Leon et al., 2009; Cohen et al., 2011; Cohen and Mizrahi, 2015; Lau et al., 2020; Carcea et al., 2021). Emergence of retrieval in adult females is accompanied by changes in the inhibitory circuitry of the auditory cortex (Liu and Schreiner, 2007; Galindo-Leon et al., 2009; Cohen et al., 2011; Lin et al., 2013; Cohen and Mizrahi, 2015; Marlin et al., 2015; Lau et al., 2020). *Mecp2* expression is specifically required in the auditory cortex at the time of pup exposure for proper retrieval (Krishnan et al., 2017). Moreover, loss of *Mecp2* triggers overexpression of parvalbumin (PV) and the extracellular matrix structures perineuronal nets (PNNs) (Krishnan et al., 2017). These two markers associated with the PVin are thought to act as ‘brakes’ on cortical plasticity (Krishnan et al., 2017). Restoration of normal levels of PV and PNN expression in the auditory cortex improved behavior and restored physiological plasticity (Krishnan et al., 2017; Lau et al., 2020). Given that changes in PVin-specific markers were correlated with retrieval behavior performance, we speculated that PVin are central to the behavioral phenotype of *Mecp2^het^*.

Loss of *Mecp2* appears to be more detrimental in certain cell types. For example, inhibitory cells may be particularly impaired by loss of *Mecp2*. Restriction of *Mecp2* mutation to either all GABAergic cells or to selected inhibitory subclasses (e.g., parvalbumin- or somatostatin-positive interneurons) is sufficient for recapitulating the majority of phenotypes in mouse models (Chao et al., 2010; He et al., 2014; Ito-Ishida et al., 2015; Mossner et al., 2020). Other work has demonstrated selected behavioral effects resulting from loss of *Mecp2* in excitatory neurons (Chao et al., 2007; Meng et al., 2016). Here we use cell type-specific removal of *Mecp2* and show that PVin are the only major class of interneurons that significantly affect retrieval when depleted of Mecp2. Mice of the genotype *PV-Cre^+^/Mecp2^flox^* (henceforth PV-Mecp2 mutants), which lack *Mecp2* in all PVin, are delayed in the onset of pup retrieval and recapitulate all the major features of *Mecp2^het^*. Specifically, when virgin PV-Mecp2 mutant females were exposed to pups, they exhibited elevated expression of PV and PNNs relative to *Mecp2^wt^*. PVin-specific mutants also did not show the experience-dependent disinhibition of auditory cortex seen in *Mecp2^wt^* controls. Finally, deleting PVin in the auditory cortex in adulthood was sufficient to delay pup retrieval. Taken together, these findings are consistent with the conclusion that *Mecp2* in auditory cortex PVin is critical for initiating experience-dependent auditory plasticity that facilitates the emergence of maternal retrieval.

## MATERIALS AND METHODS

### Animals

All procedures were conducted in accordance with the National Institutes of Health’s Guide for the Care and Use of Laboratory Animals and approved by the Cold Spring Harbor Laboratory Institutional Animal Care and Use Committee. Animals were maintained on a 12-h-12-h light-dark cycle and received food and water *ad libitum*. Behavioral experiments were conducted during light-cycle hours.

Subjects were adult, female mice 6-12 weeks of age, bred in-house from founders obtained from The Jackson Laboratory (Bar Harbor, ME) or the Mutant Mouse Resource and Research Center (Davis, CA). The following genotypes were used: CBA/CaJ, B6.129P2(C)-Mecp2^tm1.1Bird^/J (‘Mecp2^het^’, Jax #003890), B6.129S4-Mecp2^tm1Jae^/Mmucd (‘Mecp2^flox^’, MMRC #011918), B6.129P2-Pvalb^tm1(cre)Arbr^/J (‘PV-Cre’, Jax #017320), Vip^tm(cre)zjh^ /J (‘VIP-Cre’, Jax #010908), Sst^tm2.1(cre)zjh^/J (‘SS-Cre’, Jax #013044), B6.129S2^tm(emx1)krj^/J (‘Emx1-Cre’, Jax #005628), and PV-Flp B6.Cg-Pvalb^tm4.1(flpo)Hze^/J (‘PV-Flp’, Jax #022730). All crosses between Cre/Flp recombinase lines were established by pairing carriers of each allele such that all female test subject cagemates (controls and mutants) were homozygous for the *Mecp2*-flox allele and had 0 – 2 copies of the relevant recombinase allele. For example, in the PV-Mecp2 line, ‘PV-Mecp2 mutants’ were either PV-ires-Cre^+/-^ or PV-ires-Cre^+/+^ and were homozygous for Mepc2^flox^ (Mecp2^flox+/flox+^). ‘PV-Mecp2 wild type’ (WT) controls were negative for the recombinase (PV-ires-Cre^-/-^) but homozygous for Mecp2^flox^.

All animals were genotyped at the time of weaning– approximately three weeks of age– according to standard protocols from the source. In some cases, genotyping was performed by an external service (Transnetyx, Cordova, TN) using their suggested probes or Jackson Laboratory probes. All lines were monitored for the possibility of somatic recombination affecting the *Mecp2* gene as per Jackson Laboratory recommendations.

### Behavioral analysis

Pup retrieval behavior was conducted as previously described (Krishnan et al., 2017). In brief, virgin adult female test subjects (‘surrogates’) were co-housed with a wild type (CBA) pregnant dam 2-5 d pre-partum. Starting at PND 0, subjects were tested daily for 3 consecutive days in a pup retrieval assay, as follows. Pups were isolated for 2 min and then scattered to set positions in the home cage. Surrogates were allowed to interact with scattered pups for 5 min. Animals not currently performing the retrieval assay, including the dam, were temporarily placed in a group holding cage. Holding cages and home cages were not changed for the duration of retrieval experiments (i.e., from the time of surrogate pairing to PND 2). A normalized latency score between 0 (instantaneous gathering of all pups) and 1 (failure to gather all pups) was calculated as follows:

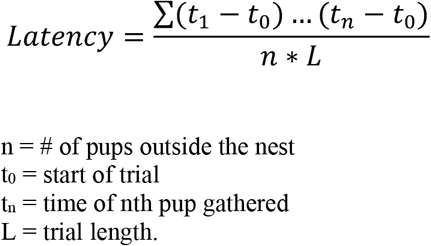

### Surgeries and injections

All surgeries were performed on a KOPF stereotaxic device. Anesthesia induction was achieved with a bolus intraperitoneal (IP) injection of ketamine (100 mg/kg) and xylazine (5 mg/kg) mixture and maintained for long surgeries (>2 h) with inhaled isoflurane (1 – 2%) in oxygen (2 – 4 Lpm), adjusted as needed based on assessment of the depth of anesthesia with a tail/paw pinch every 30 min. At the end of surgery, a non-steroidal anti-inflammatory (Meloxicam, 2 mg/kg IP) and an antibiotic (enrofloxacin, selected for its lack of ototoxicity, 4 mg/kg IP) were administered for analgesia and infection prophylaxis.

All animals used for *in vivo* physiology or fiber photometry experiments had a custom titanium headbar affixed to the skull at the time of craniotomy. To optimize the stability of the headbar, the surface of the skull was lightly, manually etched with a scalpel and three different dental cements were applied: Metabond Quick (C&B), Vitrebond Light Cure Glass Ionomer (3M), and OrthoJet (Lang). Mice used for fiber photometric recordings were also fitted with optical fiber implants secured with dental cement. Uncleaved fibers (0.39 NA, 200 μm dia., 1.25 mm length) (CFMLC12U-20, ThorLabs, Newton, NJ) were manually cleaved to the desired length using a Ruby Scribe (S90R, ThorLabs). A guide (OGL-5, ThorLabs) was also secured (Loctite, Henkel Adhesives) into place and the optical fiber implants were lowered into the auditory cortex to a depth of 700 μm. Implants were allowed to cure >48 h before head fixation or attachment of an optical cable. Braided silk surgical sutures (CP Medical) and/or Vetbond tissue adhesive (3M) were used to close surgical sites around headgear.

Thalamorecipient ‘core’ auditory cortex (Lin et al., 2013) was targeted for adeno-associated virus (AAV) injections and neurophysiology recordings using the following coordinates relative to Bregma: anterior 2.5 mm (+/- 0.4 mm) and lateral 3.9 mm (+/- 0.3 mm). For AAV-injection experiments, 90 nL injections (delivered at 20 nl/min) were administered via glass pipettes (20 μm tip) at 6 locations per hemisphere, relative to Bregma: anterior 2.1 mm, 2.4 mm, and 2.7 mm at lateral 3.8 mm and 4.0 mm. Viruses used were pAAV-EF1a-fDIO-Cre (Addgene, viral prep #121675-AAV9) or pAAV.Syn.Flex.GCaMP7s.WPRE.SV (Addgene, viral prep # 104491-AAV9).

### Immunohistochemistry

Subjects were injected with a lethal dose of Euthasol (pentobarbital sodium and phenytoin sodium cocktail, IP) and transcardially perfused with ice-cold phosphate buffered saline (PBS) followed by paraformaldehyde (PFA). Brains were extracted and fixed overnight in PFA at 4°C before being transferred to a 30% sucrose solution in PBS again at 4°C overnight or until buoyancy of the tissue was lost. Frozen 45 μm sections were collected using a sliding microtome (Leica SM 2010R). Free-floating sections were preserved for batch immunohistochemical staining (IHC) in cryoprotectant solution at −20°C. Cryoprotectant solution consisted of sucrose (0.3g/ml), polyvinyl-pyrrolidone (0.01g/ml), and ethylene glycol (0.5ml/ml) in 0.1 M PB.

For all staining protocols, free-floating sections were washed three times at room temperature (RT) in PBS and 0.3 % Triton X-100, followed with a 30 min wash in 0.3% hydrogen peroxide solution to decrease non-specific background staining. Subsequently, sections were incubated in 5% normal goat or donkey serum, in accordance with the chosen secondary antibody, and then primary antibody solution at 4°C overnight. The following primary antibodies and dilutions were used to stain for MeCP2, parvalbumin, and perineuronal nets, respectively: rabbit anti-MeCP2 (Cell Signaling Technologies, 1:1000), mouse anti-parvalbumin (Sigma, 1:1000), and lectin from Wisteria floribunda with biotin conjugate (Sigma, 1:1000). Rabbit anti-HA-Tag (Abcam, 1:500) was used on alternating sections to detect cells expressing pAAV-EF1a-fDIO-Cre in PV-Flp subjects. In a subset of animals used for fiber photometry, staining was used to amplify GCaMP expression with chicken anti-GFP (Aves, 1:1000). Sections were washed three times at RT before transfer to secondary antibody solution. Primary anybody staining was visualized with the following AlexaFluor conjugated secondary antibodies: goat AF 488, goat AF 594, and donkey AF 633 (Invitrogen Technologies). Secondary antibody dilutions matched those for the targeted primary antibody. Sections were exposed to secondary antibodies for two 2 h at RT.

### Imaging and quantification

20X magnification was used to collect z-stack images on a 710 confocal microscope (Ziess, Germany) in line scan mode. The spectra for each channel were manually adjusted to optimize signal-to-noise ratio using staining from a naïve WT sample. Those settings were as follows: bit depth = 12; laser power = 10%; gain < 700, pin hole size lowest value for each channel; full dynamic range 1024×1024 pixel smoothness, averaging = 4. These settings were used to acquire all images across batches of a given stain. For each brain, 6 – 8 matched sections spanning the rostral-caudal axis of the auditory cortex were selected for imaging. Maximum intensity projection images were generated for each field of view.

To quantify per cell staining intensity, FIJI (Image J) software was used to manually outline cells and apply area-integrated intensity in the “set measurements” panel. Background intensity readings for each section were subtracted from the cell intensity values. Cells were counted using FIJI (Image J) software and normalized to the volume of auditory cortex represented in images based on the size of the image and the depth of the z-stacks (μm x μm x μm). Mean cell density was determined by averaging counts per volume across sections.

### In vivo physiology

Loose-patch recordings from the auditory cortex were performed in awake, head-fixed subjects that were allowed to freely run on an axially rotating foam wheel as previously described (Cazakoff et al., 2014; Lau et al., 2020). Animals were habituated to head fixation and the wheel for 15 – 30 min/d for 1 – 2 d before recordings were collected. For each subject, a small craniotomy (200 μm x 200 μm) was made over the auditory cortex. Subjects were head-fixed by bolting the titanium bar implant to a frame suspended above a freely rotating foam wheel for the duration of the recording. Recordings were performed over 2 – 3 consecutive daily sessions (< 8 h). Gelfoam (absorbable gelatin sponge, Ethicon) soaked in sterile saline was used to keep the surface of the cortex moist between recording periods during the session, and craniotomy sites were covered with Kwik-Cast between days of recording.

Single unit recordings were made with a bridge amplifier (BA-03X, NPI, Tamm, Germany). Borosilicate glass micropipettes (15-40 MΩ) were pulled on a horizontal pipette puller (Model P-1000, Sutter Instruments, Novato, CA). Pipettes were filled with an intracellular solution containing in mM: 125 potassium gluconate, 10 potassium chloride, 2 magnesium chloride, and 10 HEPES. Single neurons were recorded ‘blind’ by advancing the pipette in 3 – 5 μm steps using a single-axis stepper motor and controller (Solo, Sutter Instruments) or hydraulic micromanipulator (MX610, Siskiyou Corporation) as positive pressure was applied to the tip. Brief, small injected currents (−200 pA, 200 ms) were made at 2 Hz to monitor tip resistance and the capacitance buzz feature on the amplifier was used to clear debris. Voltage signals were low pass filtered (3 kHz), digitized (10 kHz), and acquired using Spike2 software and CED hardware (Power 1401, CED, Cambridge, UK). All cells were recorded at a depth < 1 mm.

Auditory stimuli consisted of a library of 8 USVs recorded from WT CBA/CaJ mouse pups (2 – 4 d old) inside an anechoic isolation chamber (Industrial Acoustics, NY, NY) using an ultrasound microphone (Avisoft, Germany) suspended 30 cm above the pup. Stimuli were played in a pseudorandom order with a 4 s interstimulus interval. Stimulus files that had been digitally sampled at 195.3 kHz were converted to analog output via CED hardware (Power 1401). Stimuli were low pass filtered (100 kHz) and amplified with custom built hardware (Kiwa Electronics, Kasson, MN) before being output through an electrostatic speaker and driver (ED1/ES1, Tucker-Davis Technologies, Palchua, FL) 4” directly in front of the animal. Speaker output was calibrated to 65 dB SPL at the mouse’s head with a sound level meter (Extech, Model 407736) using A-weighting by comparing to an 8 kHz reference tone. The speaker had relatively flat output (±11 dB) at 4 – 100 kHz.

### Fiber photometry

PV-Cre mice (WT, *Mecp2^flox^*, and *Mecp2^het^*) were prepared by injecting a cre-dependent AAV-expressing GCaMP7s and optical fiber implants in the auditory cortex as described above. Bulk GCaMP-detected calcium signals from PVin were measured using a custom setup as described (Dvorkin and Shea, 2022). Subjects were head-fixed by bolting the titanium bar implant to a frame suspended above a freely rotating foam wheel for the duration of the recording. An optical cable (200 μm, 0.39 NA) coupled to the fiber implant was used to deliver 473 nm and 565 nm light from a pair of LEDs (LEDD1B, ThorLabs). Green emitted light was used to measure the activity-dependent fluorescence of GCaMP, while red emitted light was used to monitor and correct for potential movement or optical coupling artifacts unrelated to neural activity. No such artifacts were ever detected in our head-fixed recordings. Light from each LED was modulated at 211 Hz but 180 degrees out of phase. Before each recording session, the power of the light emitted at the tip of the patch cable was measured with a power meter (ThorLabs, PM100D) and manually adjusted to 30 – 33 μW.

Emitted light was split into separate green and red paths, bandpass filtered (Chroma Technologies, Rockingham, VT), and detected by separate photodiodes (Newport Corporation, Irvine, CA). Photodiode signals were digitally sampled at 6100 Hz via a data acquisition board (NI USB-6211, National Instruments, Austin,TX). Sinusoidal peaks of each signal were extracted by custom Matlab (Mathworks, Natick, MA) software to achieve an effective sampling rate of 211 Hz. Signals were corrected for photobleaching by fitting the decay with a double exponential function. Since head-fixed recordings were uncontaminated by artifacts, the green emission signal was used to compute ΔF/F.

The same USV stimulus set was presented during fiber photometric recordings as described for electrophysiology recordings with an interstimulus interval of 10s. Custom MATLAB software was used to present stimuli and acquire data via hardware from National Instruments.

### Experimental design and statistical analysis

All data visualization and statistical analysis was performed in Matlab or Prism (Graphpad, San Diego, CA). Unless otherwise noted, values are reported as mean ± SEM. Behavioral latency data was analyzed with a two-way ANOVA (with factors of time and genotype/treatment) and where warranted, posthoc comparisons were made. All histology was performed in batches wherein one subject from each experimental group was represented, and the scorer was blinded to the group. Per cell PV intensity was Z-scored within each batch, and for PNN counts, in each batch a threshold was applied at the mean + 2 SD for all sections in a batch. Only PNNs visible after thresholding were counted. Significant differences in PV intensity and PNN counts were statistically analyzed with a one-way ANOVA.

Spike2 software (Cambridge Electronic Design Ltd, Cambridge, UK) was used to manually threshold and sort single unit spike shapes based on PCA clustering. A total of 287 individual neurons were included in the analysis. Several previous studies, including work from our lab, have identified distinct properties of PVin waveforms, including a narrow spike shape, nearly symmetrical positive and negative peak amplitudes, and elevated firing rates (Cohen and Mizrahi, 2015; Lau et al., 2020). We combined our current data set with another 26 neuronal recordings previously collected in our lab that used photoidentification of PVin expressing the optogenetic activator ChR2. Each of the total 313 neurons was represented by a vector of 28 points defining the mean spike shape plus the cell’s ongoing firing rate. The 313 x 29 matrix was used as the input to a PCA analysis, and the result was analyzed by k-means clustering (k = 3). All optically identified neurons were contained within a single cluster; therefore, the neurons in that cluster were designated as putatively PVin.

Peristimulus time histograms (PSTHs; 10 ms bin size) were constructed of each cell’s mean firing rate in response to each stimulus (i.e. cell-stimulus pairs), and bin values were transformed to Z-scores for each cell. Significant responses were identified among all cell-stimulus pairs with a bootstrap procedure as follows. If a given stimulus was presented *n* times, *n* windows of 150 ms each were randomly chosen from the entire duration of spiking recorded for that neuron, and the mean spike rate for all *n* windows was calculated. This was repeated 10,000 times to generate a null distribution of randomized spiking rates. Significant cell-stimulus pairs were identified as those for which the actual mean response in the 150 ms after the stimulus onset fell within the upper or lower 2.5% of the spiking rate null distribution. Cells that lacked a significant response to any stimulus were discarded from the analysis. The response for each cell-stimulus pair was computed as the integrated area under the first 200 ms of the PSTH in units of Z-score*s. Mean responses across experimental groups were statistically compared with Mann-Whitney U tests.

For fiber photometry data, to compare fluorescence signals across animals and over time, ΔF/F signals collected from each animal were transformed to a Z-score. Responses to each stimulus were computed as the integrated area under the mean response curve in units of Z-score*s. For each genotype, mean responses to auditory stimuli at the pup-naïve timepoint were compared to mean responses measured at a post-naïve time point (PND 3 – 5) with a paired t-test.

## RESULTS

### Acquisition of pup retrieval is delayed by loss of Mecp2 in parvalbumin interneurons

We previously showed that female *Mecp2^het^* mice fail to reliably retrieve pups, even after 5 d of cohabitation with a WT dam and her litter (Krishnan et al., 2017). We also found that when mice were crossed between PV-Cre and *Mecp2^flox^*, PV-Mecp2 mutants were initially slower to retrieve as compared PV-Mecp2 WT subjects (Krishnan et al., 2017). This raised the possibility that certain cell types within the auditory cortex might be more important than others for the neural plasticity that facilitates retrieval. Here we replicate that finding, and we compare the results with our observations from knocking out *Mecp2* in several other genetically-restricted neuronal populations.

We used several mouse lines expressing Cre-recombinase in specific cell types in conjunction with *Mecp2^flox^* mice to restrict *Mecp2* knockout to three distinct populations of GABAergic inhibitory neurons: parvalbumin-expressing ‘PVin’, somatostatin-expressing ‘SSTin’, and vasoactive intestinal peptide-expressing ‘VIPin’. We also used the Emx1-Cre line to restrict *Mecp2* knockout to the majority (~90%) of excitatory cortical pyramidal neurons (Briata et al., 1996) (Figure 1A). Female subjects were co-housed with a pregnant WT CBA female, and beginning on PND 0, were tested daily in a pup retrieval assay (Figure 1B; see Materials and Methods) (Krishnan et al., 2017). Co-housing gives the virgin females the opportunity to observe and participate in interactions with pups (Carcea et al., 2021). Therefore, their improvement in performance over time reflects only the influence of experience, not hormonal changes related to pregnancy and parturition.

**Figure 1:**
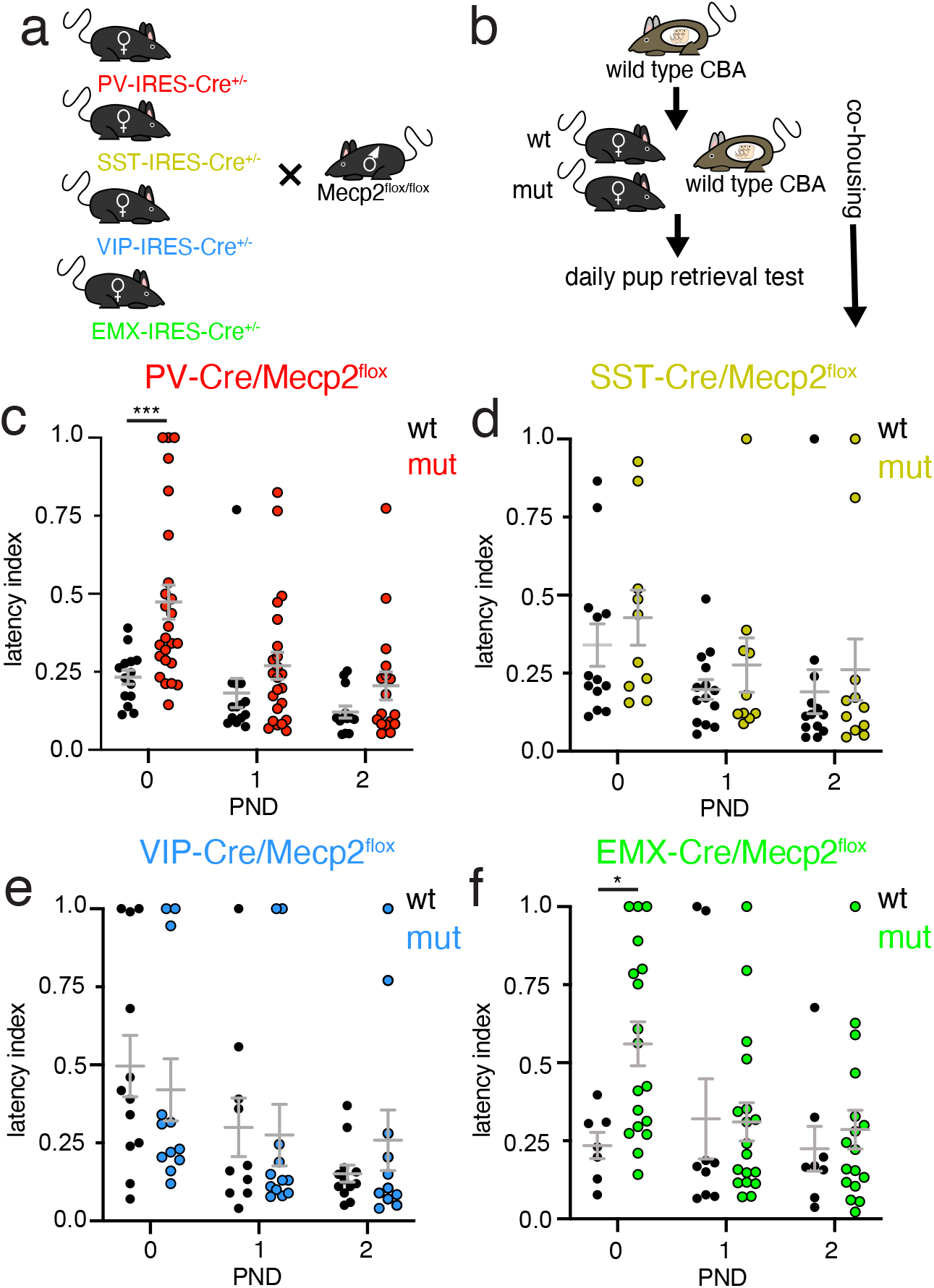
Cell-type specific deletion of *Mecp2* has varying effects on pup retrieval behavior. (A) Schematic of mouse lines crossed to achieve selective *Mecp2* deletion in different neuron types. (B) Schematic of co-housing and retrieval behavior protocol. (C – F) Scatterplots of retrieval latency comparing performance of cell type-specific Mecp2 mutants to that of their littermate controls for PV-Cre (C), SST-Cre (D), VIP-Cre (E), and Emx1-Cre (F) lines crossed with *Mecp2^flox^* mice. (C) Emergence of pup retrieval was delayed in PV-Mecp2 mutants relative to controls. A two-way ANOVA revealed significant main effects for time (day of testing), (F = 9.21, *p* < 0.001), and genotype, (F = 9.41, *p* < 0.01), but not an interaction (F = 1.94, *p* = 0.15). Post hoc tests revealed a significant difference between mutant and WT animals on PND 0 only (n = 14 controls, latency: 0.17 ± 0.03; n = 25 mutants, latency: 0.32 ± 0.08; Sidak’s test, \*\*\**p* < 0.001). (D) Timing of emergence of pup retrieval did not differ between SST-Mecp2 mutants and controls. A twoway ANOVA revealed a significant main effect for time (day of testing), (F = 8.98, *p* < 0.01), but not for genotype or for an interaction. Post hoc tests revealed no significant difference between mutant and WT animals on any day (Sidak’s test, *p* > 0.05). (E) Emergence of pup retrieval did not differ between VIP-Mecp2 mutants and controls. A Two-way ANOVA revealed a significant main effect for time/day (VIP-Mecp2: F = 7.50, *p* < 0.01), but not for genotype, nor an interaction between those variables. Post hoc tests revealed no significant difference between mutant and WT animals on any day of testing (Sidak’s test, *p* > 0.05). (F) Emergence of pup retrieval was delayed in Emx1-Mecp2 mutants relative to controls. A two-way ANOVA revealed a significant effect only for the interaction between time and genotype, (F = 3.86, *p* < 0.05), but neither a main effect for genotype, (F = 2.45, *p* = 0.13), nor day, (F = 2.13, *p* = 0.14). A post hoc test revealed a significant difference between mutant and WT animals for PND 0 only (n = 7 controls, latency: 0.26 ± 0.03; n = 18 mutants, latency: 0.39 ± 0.09; Sidak’s test, \**p* < 0.05).

PV-Mecp2 mutants showed significantly longer retrieval latency scores compared to PV-Mecp2 WT on PND 0. However, these subjects improved over time, matching the performance of PV-Mecp2 WT mice by PND 1 (Figure 1C). A two-way mixed effects ANOVA revealed significant effects of time (day) (F = 9.21, *p* < 0.001) and genotype (F = 9.41, *p* < 0.01), but not an interaction between those variables (F = 1.94, *p* = 0.15). Posthoc testing revealed a significant difference between the mutant and WT groups for PND 0 only (Sidak’s test, *p* < 0.001). Individual unpaired comparisons for each day detected a significant difference between genotypes only on PND 0 (n = 25 mutant, 15 WT; Mann-Whitney corrected for multiple comparisons, *p* < 0.01). Therefore, PV-Mecp2 mutants showed a transient disruption in pup retrieval.

To assess the specificity of this result to the PVin population, as opposed to other interneuron types, we ran the same experiment with mice in which *Mecp2* was knocked out in one of two other major classes of GABAergic inhibitory neurons (SSTin or VIPin) (Figure 1D, E). Two-way mixed effects ANOVAs revealed significant effects for time (day) in both cohorts (SST-Mecp2: F = 8.98, *p* < 0.01; VIP-Mecp2: F = 7.50, *p* < 0.01), but not for genotype, or for an interaction between those variables. Post hoc testing revealed no difference between mutant and WT groups for any day of testing (Sidak’s test, *p* > 0.05). Individual unpaired comparisons for each day also failed to detect significant differences between genotypes on any day (*p* > 0.05) for either line. Therefore, neither SST-Mecp2 mutants nor VIP-Mecp2 mutants showed a transient disruption in pup retrieval as observed for PV-Mecp2 subjects.

As a comparison to *Mecp2* deletion in small interneuron populations, we next crossed *Mecp2^flox^* mice with the Emx1-Cre line to restrict *Mecp2* knockout to roughly 90% of excitatory neurons in the cortex and hippocampus (Briata et al., 1996). Like PV-Mecp2 mutants, Emx1-Mecp2 mutants exhibited a delayed onset of pup retrieval (Figure 1F). A two-way mixed effects ANOVA revealed a significant effect only for an interaction (F = 3.86, *p* < 0.05), but neither a main effect for genotype (F = 2.45, *p* < 0.13), nor for day (F = 2.13, *p* = 0.14). Post hoc testing revealed a significant difference between mutant and WT groups for PND 0 only (Sidak’s test, *p* < 0.01). Individual unpaired comparisons for each day detected a significant difference between genotypes only on PND 0 (n = 18 mutant, 7 WT; Mann-Whitney corrected for multiple comparisons, *p* < 0.05). Therefore, like PV-Mecp2 mutants, Emx-Mecp2 mutants showed a transient disruption in pup retrieval. Interestingly, this disruption was comparable in the two groups, despite the disparity in the size of the cell populations.

### Loss of Mecp2 only in PVin recapitulates changes to inhibitory markers in Mecp2^het^

High levels of expression of PVin markers (parvalbumin ‘PV’ and perineuronal nets ‘PNNs’) are taken as an indicator of maturity in PVin and are well-correlated with reduced capacity for synaptic plasticity and learning in development and adulthood (Pizzorusso et al., 2002; Carulli et al., 2010; de Vivo et al., 2013; Donato et al., 2013; Happel et al., 2014; Hou et al., 2017; Cisneros-Franco and de Villers-Sidani, 2019; reviewed in Rupert and Shea, 2022). Previously, we reported that both markers exhibited overexpression in the auditory cortex of *Mecp2^het^* after 5 d of exposure of a virgin female to pups (Krishnan et al., 2017). This experience-dependent overexpression was not observed in *Mecp2^wt^*, and genetic and pharmacological approaches that reversed it restored retrieval performance in *Mecp2^het^* (Krishnan et al., 2017).

Given that *Mecp2* deletion in PVin, a small population of neurons (~10%), is sufficient to disrupt retrieval behavior (albeit temporarily), and that population is also the locus of key pathological features in *Mecp2^het^* models, we hypothesized that the changes to PV and PNN may reflect a cell-autonomous consequence of *Mecp2* deletion from PVin. To test this hypothesis, we compared the level of PV protein and PNN expression by auditory cortex PVin between PV-Mecp2 mutants and PV-Mecp2 WTs. We did this by performing IHC and confocal microscopy of fixed brain sections from naïve mice (pre pup exposure), and experienced mice (post pup exposure) at the PND 1 and PND 5 time points (Figure 2A, B). We quantified per cell intensity of PV staining and, to minimize batch effects, converted intensities from each batch to a Z-score. We compared the distribution of Z-scores for each group, focusing on the changes within each genotype across timepoints as pup experience increased. A one-way ANOVA revealed significant differences among group means (F = 47.3, *p* < 0.001). PV-Mecp2 WT virgin mice showed a drop in PV expression; PV staining intensity was significantly lower on PND 5 compared to intensities of the naïve and PND 1 cohorts (Figure 2C, Sidak’s test, *p* < 0.001). In contrast, PV-Mecp2 mutants exhibited an increase in PV expression; PV staining intensity was significantly higher in tissue collected at both PND 1 and PND 5, as compared to naïve animals (Sidak’s test, *p* < 0.001)

**Figure 2:**
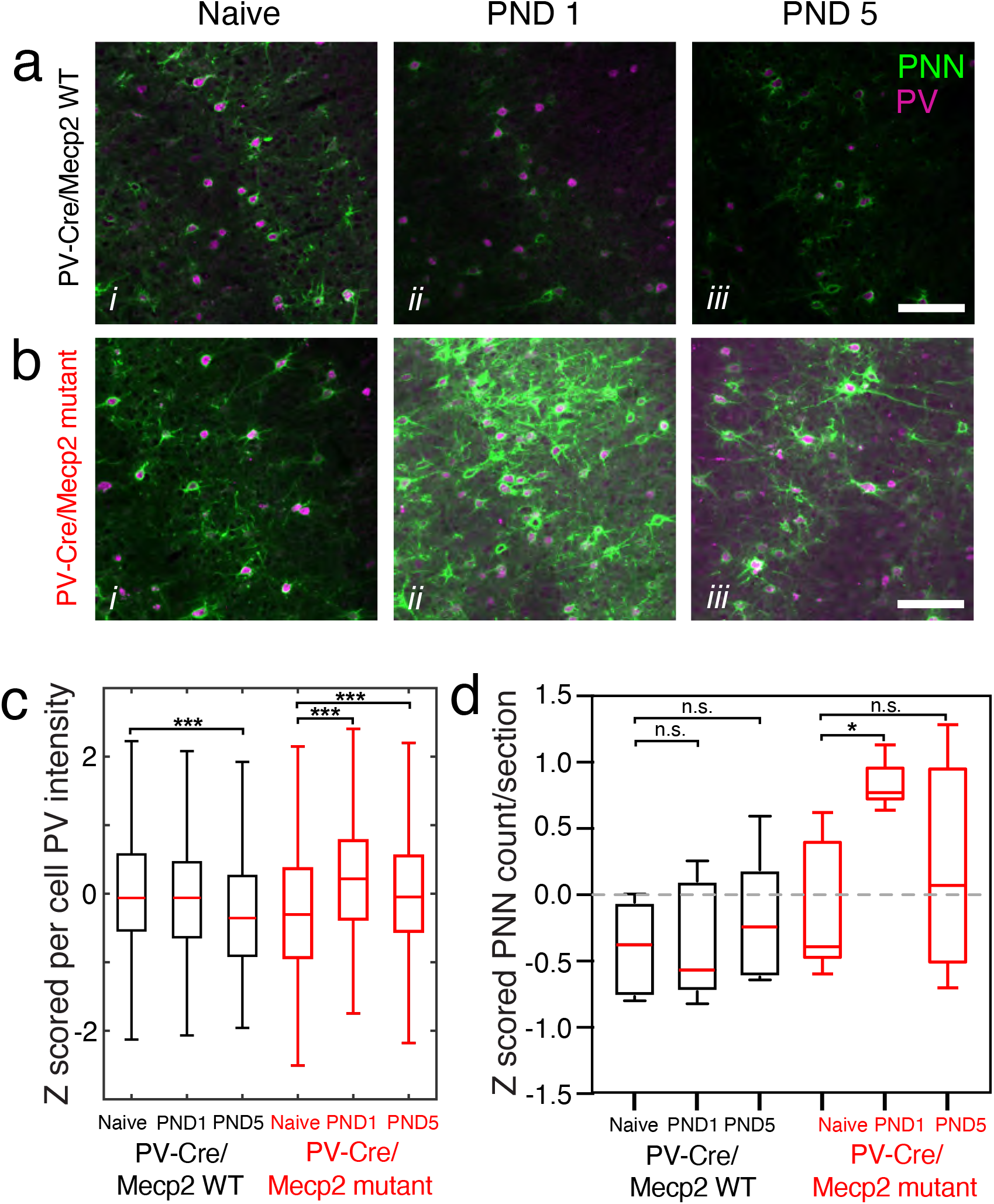
Selective deletion of *Mecp2* in PVin recapitulates changes to inhibitory markers in *Mecp2^het^*. (A) Series of photomicrographs of example sections of the auditory cortex taken from a PV-Mecp2 WT naïve mouse (i), a mouse after 2 d of cohabitation (PND 1) (ii), and a mouse after 6 d of cohabitation (PND 5) (iii). All sections were stained via IHC for parvalbumin (purple) and PNNs (green). Scale bar = 200 mm. (B) Same as (A) but for sections taken from PV-Mecp2 mutants. (C) Boxplot of the distributions of per-cell intensity of parvalbumin staining. All histology was run in six batches, with one brain from each genotype-condition group represented in each batch. All individual neuron intensities were Z-scored per batch. The total number of neurons in each group is (left to right): 1006, 996, 757, 1108, 1084, 1063. A one-way ANOVA detected differences among the groups (F = 47.3, *p* < 0.001). Post hoc testing revealed a significant decrease of mean PV intensity in PV-Mecp2 WT on PND 5 compared to naïve mice (naïve latency: 0.076 ± 0.03 Z-score; PND 5 latency: 0.25 ± 0.03 Z-score; Tukey’s test, *p* < ***0.001). In contrast, PV-Mecp2 mutant mice had higher mean per cell intensity PV staining on PND 1 (0.31 ±0.03 Z score) and PND 5 (0.047 ± 0.03 Z-score) as compared to naïve mice (−0.25 ± 0.03 Z-score; Tukey’s test, *p* < ***0.001). (D) Box plot of mean high intensity PNNs per section, comparing all six groups of mice. One mouse from each group was processed in each batch with total of 5 batches. Persection counts were Z-scored for all sections in each batch. (E) Considering all groups, there was a significant difference among them (one-way ANOVA, F = 4.3, *p* < 0.01). Post hoc comparisons of each experienced time point to the naïve time point for each genotype showed that PV-Mecp2 mutant mice on PND 1 had significantly more high-intensity PNNs per section than naïve PV-Mecp2 mutant mice (n = 5 mice/group; naïve mutant: −0.11 ± 0.23 Z score; mutant PND 1: 0.83 ± 0.08 Z score; Sidak’s test, \**p* < 0.05).

To quantify changes in PNN expression, we counted high intensity PNNs in the auditory cortex in the same set of sections analyzed above (see Materials and Methods). To minimize batch effects, all images were thresholded and binarized at 2 SD above the mean pixel value for each staining batch, and the counts of PNNs per section were Z-scored within each batch. An analysis of all groups detected significant differences among the groups (one-way ANOVA; F = 4.30, *p* < 0.01) (Figure 2D). Post hoc tests comparing PNN counts from experienced mice at PND 1 and PND 5 to counts at the naïve timepoint for both genotypes showed PNN counts per section was only significantly higher on PND 1 in PV-Mecp2 mutants (Sidak’s test, *p* < 0.05). In light of all these observations, we conclude that deletion of *Mecp2* in PVin is sufficient to at least transiently evoke overexpression of molecular markers closely associated with suppression of plasticity upon exposure to pups.

### PV-Mecp2 mutants lack the auditory cortical disinhibition triggered by pup exposure in WT

Our next goal was to determine whether deletion of *Mecp2* only in PVin was sufficient to reproduce the neurophysiological changes we observed in the auditory cortex of pup-experienced *Mecp2^het^* mice. We found that pup-experienced *Mecp2^wt^* mice exhibited a dramatic decrease in spiking output by auditory cortex PVin relative to that from naïve females (Lau et al., 2020). Moreover, we discovered that this disinhibition of the auditory cortex by PVin was absent in *Mecp2^het^* (Lau et al., 2020). We hypothesized that deletion of *Mecp2* only from PVin may affect their stimulus-evoked firing in a cell-autonomous manner. To test this hypothesis, we made loose-patch, single-unit electrophysiological recordings from auditory cortical neurons in awake head-fixed animals of both genotypes at naïve and post-pup experienced timepoints. We made neuronal recordings from four experimental groups of mice: PV-Mecp2 mutant mice that were naïve to pups (‘PV-Cre/Mecp2 mutant Naive’; n = 7 mice), PV-Mecp2 mutant mice that co-habitated with a WT dam and her pups for >5 d (‘PV-Cre/Mecp2 mutant Experienced’; n = 10 mice), PV-Cre/Mecp2 WT control littermates without pup experience (‘PV-Cre/Mecp2 WT Naïve’; n = 15 mice), and WT littermates that experienced co-habitation (‘PV-Cre/Mecp2 WT Experienced’; n = 21 mice).

As previously reported (Wu et al., 2008; Oswald and Reyes, 2011; Cohen and Mizrahi, 2015; Lau et al., 2020), PVin and non-PV neurons had characteristic spike shapes that could be distinguished by their features. We therefore combined the neurons we recorded here with a wild-type data set from a previous study (Lau et al., 2020) in which we optically identified ChR2-expressing PVin. We identified putative clusters of PVin and non-PV neurons in a principal components analysis (PCA) with a K-means clustering algorithm (see Materials and Methods). Average spike waveforms for our putatively identified populations of cells are plotted as corresponding color traces in Figure 3B and C. PVin had particularly narrow spike waveforms and were more symmetrical in amplitude around the baseline; non-PV neurons were wider and had more prominent positive peaks (Figure 3B, C). Despite our use of a novel PCA-based sorting method, our results were very consistent with previous classification results from our group and others (Wu et al., 2008; Oswald and Reyes, 2011; Cohen and Mizrahi, 2015; Lau et al., 2020).

**Figure 3:**
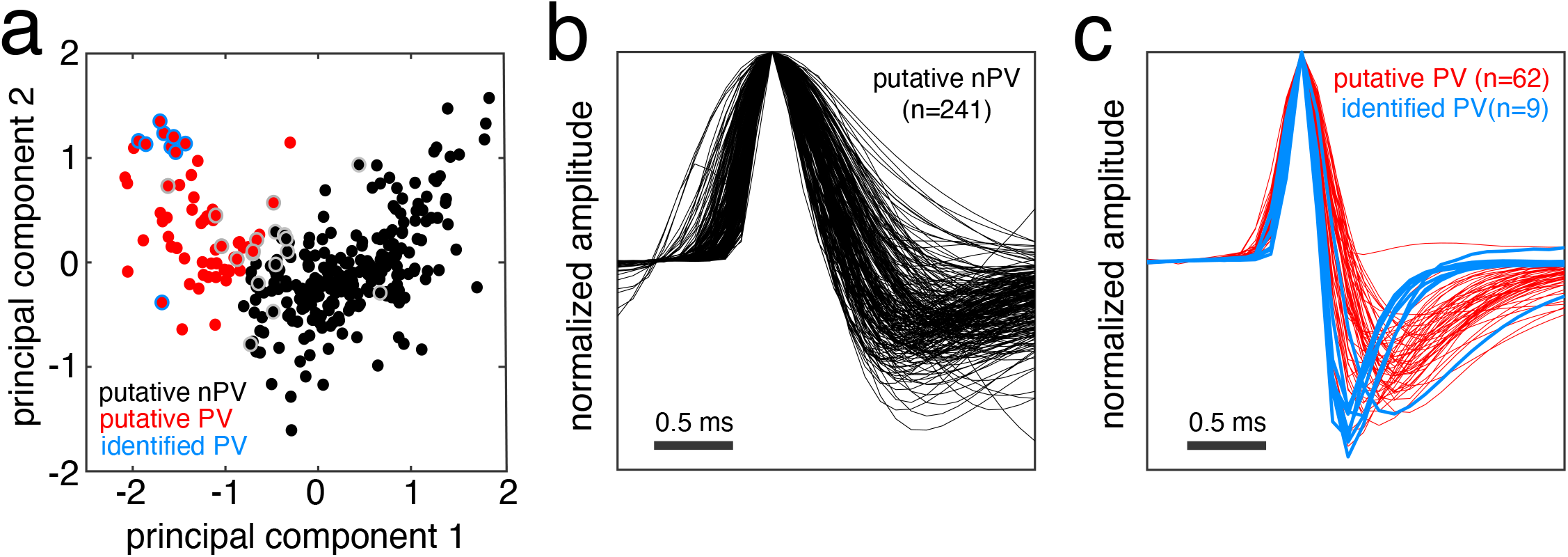
Classification of PVin and non-PVin single unit recordings by spike waveform and firing rate. (A) Scatterplot showing the results of a principal components analysis and k-means clustering analysis of 313 auditory cortex neurons based on the mean spike waveform and baseline firing rate. Red points denote putative PVin, black points denote putative non-PV neurons, points with blue border denote subset of optically-tagged PVin, and points with gray border denote inconclusive optical identification results. (B) Plot of all mean waveforms from putative non-PV neurons (n=241). (C) Plot of all mean waveforms from putatively identified PVin (n=71).

We first examined the responses of non-PV neurons from each group to a library of 8 USVs recorded from pups that were 2 – 4 d old (Lau et al., 2020). We observed that individual non-PV neurons often exhibited distinct responses to different USVs, responding with either increases or decreases in firing. Therefore, we identified all cell-call pairs (mean responses of one neuron to one stimulus) that exhibited a statistically significant change in firing rate as assessed with a bootstrap procedure (see Materials and Methods). A peristimulus time histogram (PSTH; bin size = 10 ms) was constructed to visualize the mean response of each cell to each stimulus, and the bins of all PSTHs from each cell were transformed to a Z-score.

Heatmaps in Figure 4A and B depict 2-dimensional PSTHs that each represent the mean responses for all cell-call pairs from one of the four groups of wild types (Figure 4A) and mutants (Figure 4B). Rows in each 2d-PSTH are sorted from the largest firing decrease to the largest firing increase measured in the 200 ms window after stimulus onset. We separately compared the mean of all ‘excitatory’ responses (stimulus-driven increase in firing rate) and the mean of all ‘inhibitory’ responses (stimulus-driven decrease in firing rate) between naïve and pup-experienced groups for each genotype. Figure 4C and D depict mean ± SEM traces for each sign of response. Gray traces represent recordings collected from naïve mice and purple traces represent recordings collected from pup-experienced mice. We integrated the area under the curve (AUC) for each cellcall pair and compared the distribution of response magnitudes between naïve and experienced mice cohorts. Figure 4E summarizes the results of these comparisons for PV-Mecp2 and WT mice. In WT mice, mean inhibitory responses were significantly weaker in mice that had co-housing experience with pups relative to mice that lacked pup exposure (n = 54 naïve and 44 experienced cell-call pairs; Mann-Whitney U test, *p* < 0.01). Mean excitatory responses were unchanged between naïve and experienced mice (n = 108 naïve and 72 experienced cell-call pairs; Mann-Whitney U test, *p* = 0.09). Figure 4F shows the corresponding results for PV-Mecp2 mutant mice. In these mice, mean inhibitory responses were significantly stronger in pup experienced mice than they were in pup naive mice (n = 31 naïve and 30 experienced cell-call pairs; Mann-Whitney U test, *p* < 0.001). As in WT mice, mean excitatory responses were unchanged between naïve and experienced PV-Mecp2 mutant mice (n = 60 naïve and 45 experienced cell-call pairs; Mann-Whitney U test, *p* = 0.61).

**Figure 4:**
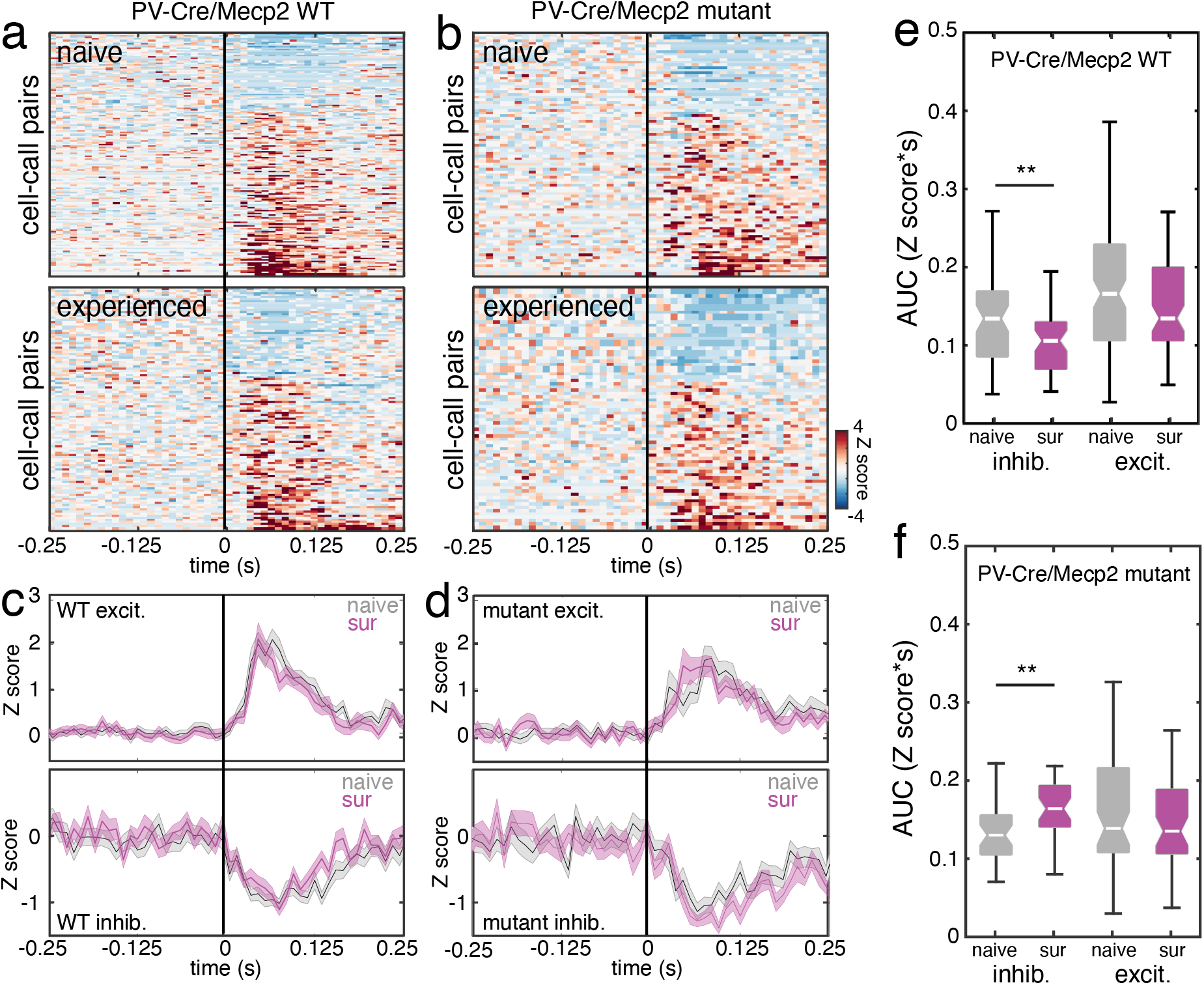
Selective deletion of Mecp2 in PVin recapitulates the suppression of non-PV neuron inhibitory plasticity seen in Mecp2^*het*^. (A) 2d-PSTHs representing the mean responses for all non-PV cell-call pairs with significant responses to USVs from naïve (top) and experienced (bottom) PV-Mecp2 WT mice. Each row within the PSTHs represents the Z-scored response of one cell-call pair. Rows are sorted from the greatest stimulus-evoked decrease in firing rate to the greatest stimulus-evoked increase firing rate. Response window of 200 ms after the stimulus onset is marked by the vertical black line. (B) Same as (A), but data are taken from auditory cortex recordings in PV-Mecp2 mutant mice. (C) Mean ± SEM traces for ‘excitatory’ responses (top) and ‘inhibitory’ responses (bottom), comparing data from naïve (gray) and experienced (purple) PV-Mecp2 WT mice. (D) Same as (C), but the data are from PV-Mecp2 mutant mice. (E) Box plot of the integrated area under the curve (AUC) for each non-PV cell-call pair recorded from PV-Mecp2 WT mice comparing the distribution of inhibitory and excitatory response magnitudes between naïve mice and experienced mice. Mean inhibitory responses were significantly weaker in pup-experienced mice relative to pup-naïve mice (naïve: n = 54 cell-call pairs, 0.132 ± 0.01 Z-score*s; experienced: n = 44 cell-call pairs, 0.101 ± 0.01 Z-score*s; Mann-Whitney U test, \*\**p* < 0.01). Mean excitatory responses were unchanged between naïve and experienced mice (naïve: n = 108 cell-call pairs, 0.195 ± 0.01 Z-score*s; experienced: n = 72 cell-call pairs, 0.161 ± 0.01 Z-score*s; Mann-Whitney U test, *p* = 0.09). (F) Same as (E), but data are from non-PV cell-call pairs recorded from PV-Mecp2 mutant mice. Mean auditory cortex inhibitory responses were significantly stronger in experienced mice than they were in pup-naive mice (naïve: n = 31 cell-call pairs, 0.134 ± 0.01; experienced: n = 30 cell-call pairs, 0.170 ± 0.01; Mann-Whitney U test, \*\*\**p* < 0.001). As in WT mice, mean excitatory responses were unchanged between naïve and experienced PV-Mecp2 mutants (naïve: n = 60 cell-call pairs 0.171 ± 0.01, Z-score*s; experienced: n = 45 cell-call pairs, 0.160 ± 0.01 Z-score*s; Mann-Whitney U test, *p* = 0.61).

We performed a similar analysis on recordings collected from putative PVin (Figure 5). In this case, because all USV responses we observed in PVin evoked increased spiking, we included all responses to stimuli for all neurons that had a significant response to at least one USV. As in Figure 4, a PSTH for each cell-call pair was constructed and organized into a 2d-PSTH for each group where rows were sorted from the weakest response to the strongest response (Figure 5A, B). Traces of mean ± SEM firing rate across all PVin cell-call pairs are plotted for naïve mice (gray) and experienced mice (purple) of each genotype (Figure 5C, D). Consistent with our previous report (Lau et al., 2020), auditory cortex PVin in experienced WT mice exhibited dramatically and significantly weaker responses to USVs compared with PVin in naïve WT mice (Figure 5B; n = 80 naïve and 104 experienced cell-call pairs; Mann-Whitney U test, *p* < 0.001). In contrast, mean responses of PVin were unchanged between naïve and pup-experienced PV-Mecp2 mutants (n = 64 naïve and 32 experienced cell-call pairs; Mann-Whitney U test, *p* = 0.61). Based on all the data from our electrophysiology experiments, we conclude that selective deletion of *Mecp2* in PVin is sufficient for disrupting the auditory cortical disinhibition that is triggered by exposure to pups as observed in WT virgin mice.

**Figure 5:**
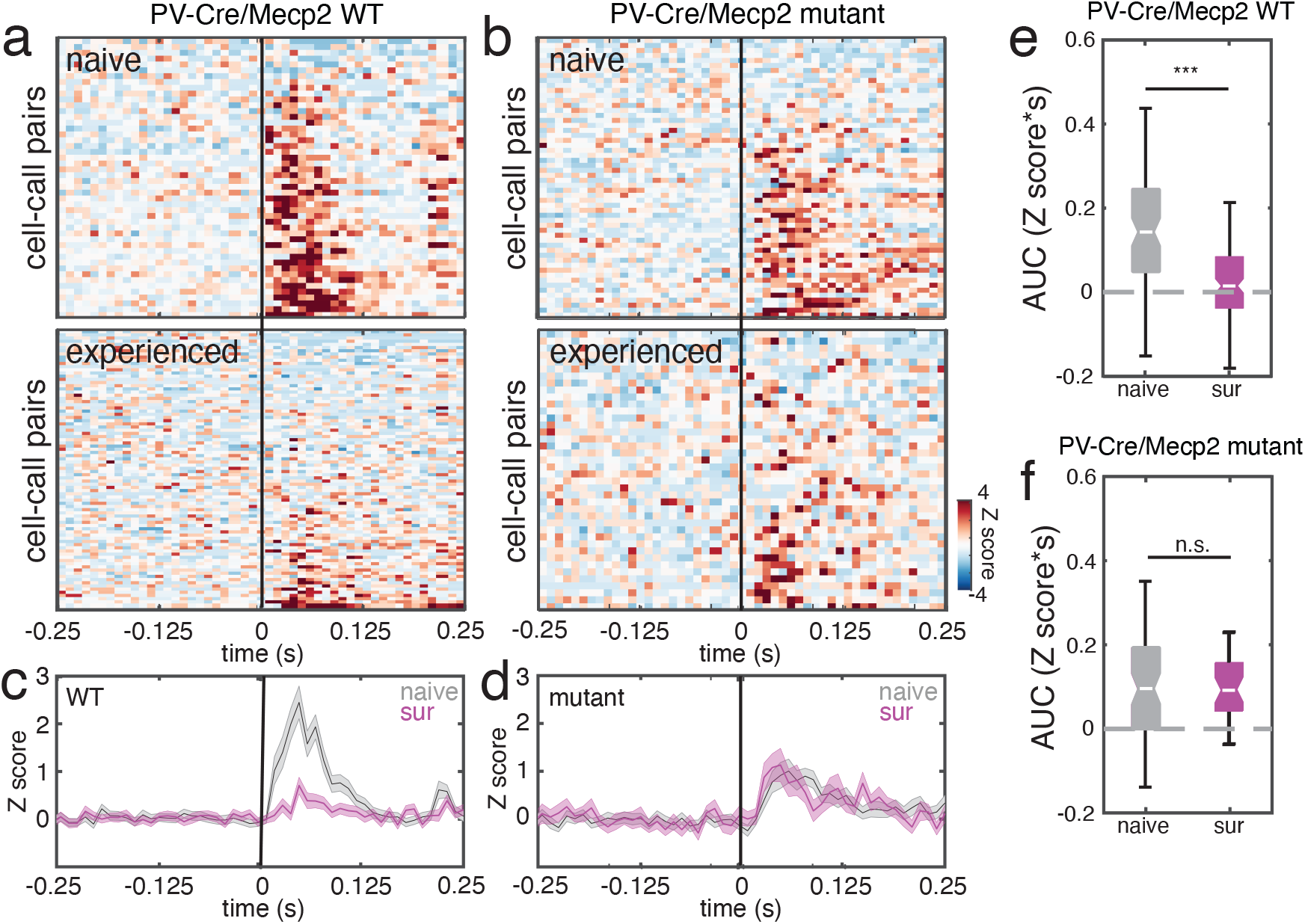
Selective deletion of Mecp2 in PVin recapitulates the suppression of PVin neuron inhibitory plasticity seen in Mecp2^*het*^. (A) 2d-PSTHs representing the mean responses for all PVin cell-call pairs with significant responses to any USV from naïve (top) and experienced (bottom) PV-Mecp2 WT mice. Each row within the PSTHs represents the Z-scored response of one cell-call pair. Rows are sorted from the greatest stimulus-evoked decrease in firing rate to the greatest stimulus-evoked increase firing rate. Response window of 200 ms after the stimulus onset is marked by the vertical black line. (B) Same as (A), but the data are from PV-Mecp2 mutant mice. (C) Mean ± SEM traces for USV responses of all cell-call pairs, comparing data from pup-naïve (gray) and pup-experienced (purple) WT mice. (D) Same as (C), but the data are from PV-Mecp2 mutant mice. (E) Box plot of the integrated area under the curve (AUC) for each PVin cellcall pair in PV-Mecp2 WT mice comparing response magnitudes between naïve mice and experienced mice. Mean responses were significantly weaker in mice that had experience with pups relative to pup-naïve mice (naïve: n = 80 cell-call pairs, 0.147 ± 0.03 Z-score*s; experienced: n = 104 cell-call pairs, 0.035 ± 0.01 Z-score*s; Mann-Whitney U test, \*\*\**p* = 0.001). (F) Same as (E), but data are from PVin cell-call pairs collected from PV-Mecp2 mutant mice. In contrast to WT mice, PVin responses were unchanged between naïve and experienced PV-Mecp2 mutant mice (naïve: n = 64 cell-call pairs 0.087 ± 0.01 Z-score*s; experienced: n = 32 cell-call pairs, 0.095 ± 0.01 Z-score*s; Mann-Whitney U test, *p* = 0.79).

### Optical recordings reveal that PVin disinhibition depends on experience and Mecp2 in PVin

One important limitation of our electrophysiology data is that it was not practical to conduct recordings from the same subjects in both naïve and pup-experienced states. A more powerful experimental design would be to measure PVin activity in the same animal over the duration of its co-habitation experience with pups. A second limitation is that these recordings yield information about only one PVin at a time. It would be useful to complement those data with recordings of the neuronal population. Therefore, we employed fiber photometry to make longitudinal measurements of widespread PVin activity in response to auditory stimuli as mice advanced from the pup-naïve state through several days of cohabitation. Subjects were prepared by making injections of Cre-dependent AAV-DIO-GCaMP7s into the auditory cortex and by implanting an optical fiber at the same location (see Materials and Methods). Because PV-Cre was necessary in all mice to express GCaMP, subjects were either *Mecp2^flox+/flox+^* (PV-Mecp2 mutant) or *Mecp2^flox-/flox-^* (PV-Mecp2 WT). We also included a group of mice that were PV-Cre+ and *Mecp2^het^* to compare results between PVin-selective *Mecp2* deletion to the non-conditional, mosaic model. We conducted daily recording sessions from head-fixed mice, presenting the same set of USVs used in the neuronal recordings.

Figure 6A shows example data for three different mice. The top row of plots represents data from a PV-Mecp2 WT subject. Each row in the heatmaps is a single trial response to one specific USV. Trials above the green horizontal line were taken from sessions prior to cohabitation (naïve state) and trials below the line were taken from sessions on PND 3 – PND 5. Each call, denoted by the black vertical tick mark, elicited an abrupt increase in fluorescence that decayed over the course of 2 s. Below each heatmap is a trace of the mean ± SEM calcium response from all naïve trials (gray) and trials collected on day 3 – 5 of pup experience (purple).

**Figure 6:**
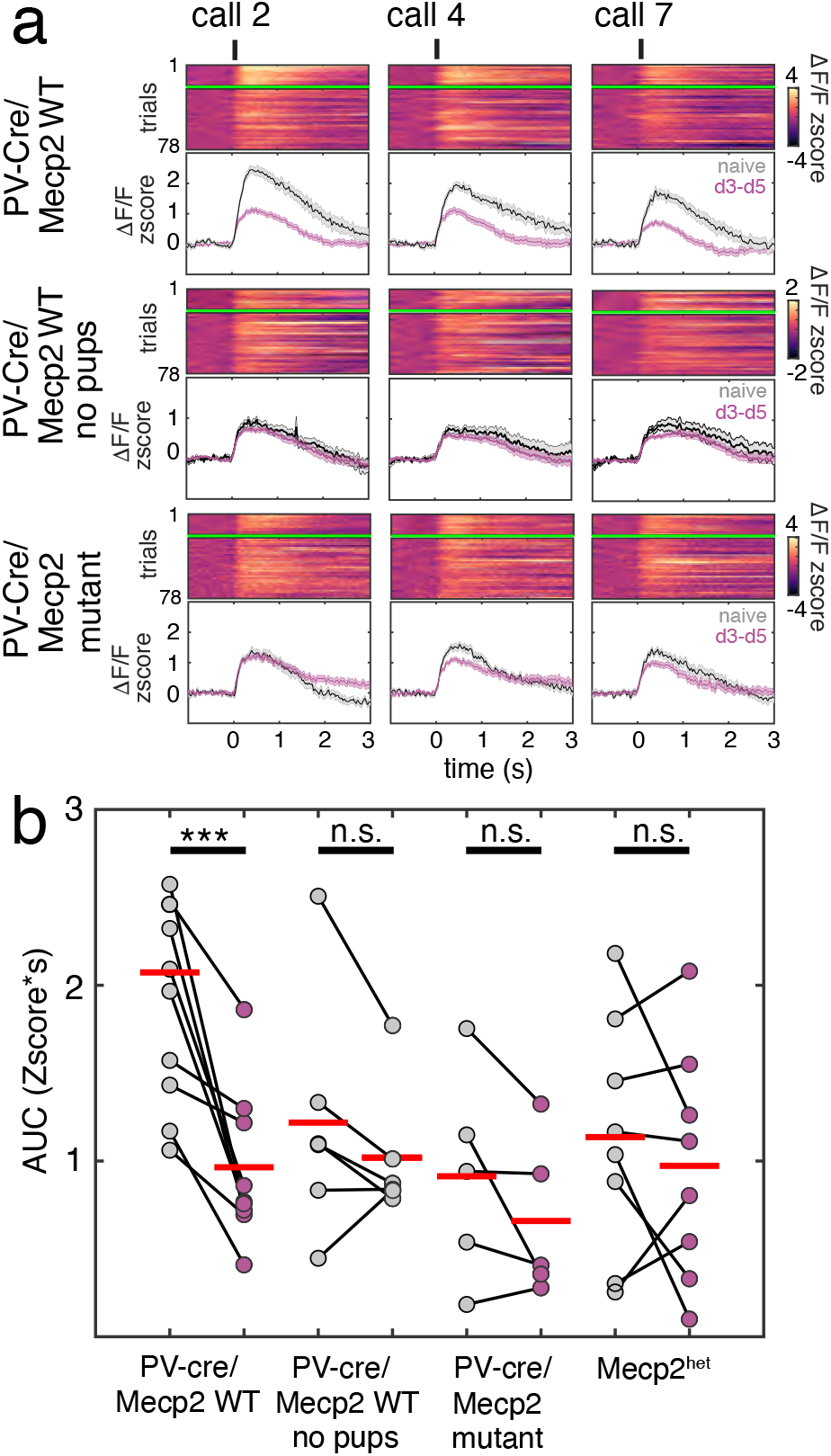
PVin disinhibition is widespread and depends upon experience and presence of Mecp2 in PVin. (A) Comparison of longitudinal fiber photometry data from three sample subjects. Fluctuations in bulk fluorescence were measured using GCaMP7s expressed in auditory cortical PVin. Each column shows the responses of each mouse to a different USV call exemplar. The top row depicts data from a PV-Mecp2 WT mouse. The middle row depicts data from a PV-Mecp2 WT mouse that was never introduced to or co-housed with pups. The bottom row depicts data from a PV-Mecp2 mutant mouse. The heatmaps depict the response to each USV over many trials gathered over several days. Each heatmap row is one trial; those above the green line were taken from sessions before pup exposure (naïve timepoint), and those below the line were taken from sessions on PND 3 – PND 5. Below each heatmap is a plot of mean ± SEM fluorescence traces from naïve (gray) and experienced (purple) timepoints. The onset of call playback is marked with a black tick above the heatmap. (B) Scatterplot of mean naïve and experienced PVin responses to all USVs for all mice in each experimental condition. Responses were quantified as the AUC of the Z-scored fluorescence trace during the first 2 s after stimulus onset. WT mice showed a consistent and significant decrease in response strength to USV between naïve trials and that during trials on days 3 – 5 of pup experience (n = 9 mice; naïve: 2.46 ± 0.79 Z-score*s; experienced: 1.03 ± 0.57 Z-score*s; paired t-test, *p* ***< 0.001). No significant differences between the early time point and the late time point responses were found for WT mice that were not exposed to pups but which were imaged during USV playback on the same schedule (n = 6 mice; naïve: 1.57 ± 1.0 Z-score*s; experienced: 1.24 ± 0.61 Z-score*s; paired t-test, *p* = 0.14), PV-Mecp2 mutant mice (n = 5 mice; naïve: 1.61 ± 1.2 Z-score*s; experienced: 1.29 ± 0.68 Z-score*s; paired t-test, *p* = 0.23), or Mecp2^*het*^ (n = 8; naïve:1.24 ± 0.77 Z-score*s; experienced: 1.11 ± 0.83 Z-score*s; paired t-test, *p* = 0.62).

We quantified the responses to calls as the mean AUC across trials and stimuli and compared that measure for all mice before and after pup exposure (Figure 6B). In agreement with our single neuron data, we found that the auditory cortex PVin population in PV-Mecp2 WT subjects exhibited consistently weaker mean responses to USVs after several days of pup exposure as compared to responses measured in the naïve state (Figure 6B; n = 9 mice; paired t test with Bonferroni correction, *p* < 0.001). Importantly, this drop in PVin responses required experience; pup-naïve control mice who were recorded on the same schedule, did not show a significant decrease in PVin activity in response to the same stimulus set presented to the experienced mice (Figure 6B; n = 6 mice; paired t test, *p* = 0.40). Neither *Mecp2^het^* (n = 8 mice; paired t test, *p* = 0.16) nor PV-Mecp2 mutants (n =5 mice; paired t test, *p* = 0.56) showed a significant decrease in the responses of PVin to USVs.

### Acquisition of pup retrieval is delayed by adult loss of Mecp2 in auditory cortical PVin

All of the above observations indicate that PVin play an important early role in initiating cortical plasticity in response to sensory and social experience with pups. Loss of *Mecp2* exclusively in this neuronal subtype, which represents only about 10% of the neurons in the neocortex, replicates many key features of the maternal behavioral and neural pathology seen in *Mecp2^het^*. However, our approach of crossing PV-Cre mice with *Mecp2^flox^* does not specifically implicate PVin in the auditory cortex, nor does it distinguish between an acute requirement for *Mecp2* in PVin in adulthood (e.g., during the virgin mouse’s initial exposure to pups) and an earlier requirement for *Mecp2* in PVin for proper development of the auditory cortex to support later plasticity. We therefore devised an intersectional viral-genetic strategy to address this limitation.

Figure 7A is a schematic depiction of our strategy to target Mecp2 only in PVin in the auditory cortex, and only after cortical development (see Materials and Methods for details). Briefly, we crossed *Mecp2^flox^* mice with a line that expresses Flp recombinase in PV neurons, and then at 6 weeks of age, bilaterally injected the auditory cortex with either an AAV driving the Flp-dependent expression of Cre recombinase and an HA tag (‘Flp-Flox’) or a control vector expressing GFP (‘GFP-Flox’).

**Figure 7:**
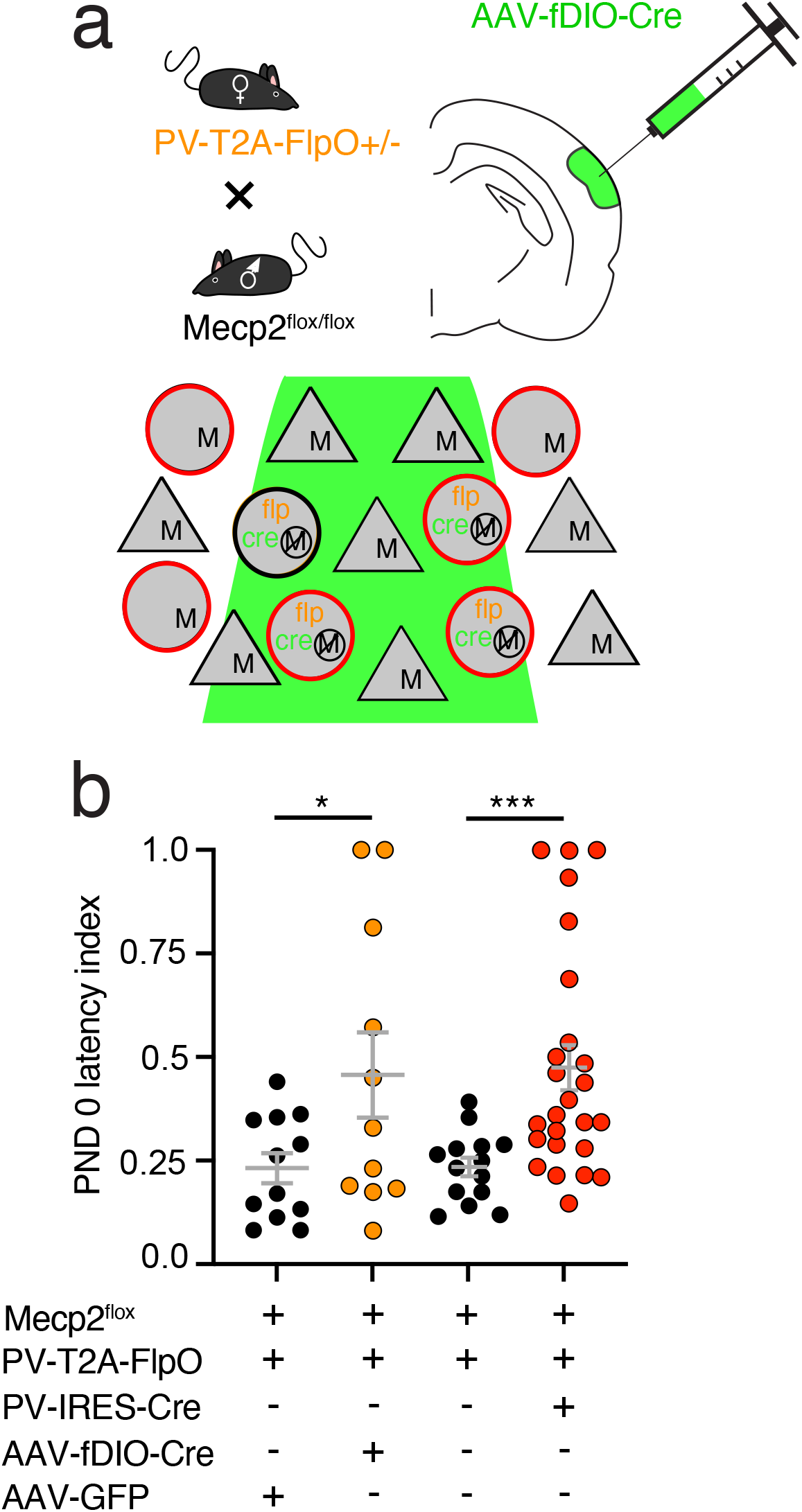
Acquisition of pup retrieval is delayed by adult loss of Mecp2 in auditory cortex PVin. (A) Schematic depiction of our experimental strategy. Mice carrying Flp recombinase after a T2A site in PV neurons were crossed with Mecp2*^flox^* mice. Offspring mice positive for both alleles were injected with an AAV driving the expression of either Flp-dependent (fDIO) Cre or GFP. The consequence of injecting fDIO-Cre is the deletion of *Mecp2* from PV neurons at the time and location of our choosing – in this case, the auditory cortex of young adult mice, thereby deleting *Mecp2* in PVin at the injection site. (B) Swarm plot comparing retrieval latencies for control subjects that were injected with AAV-GFP (left black points) to those for subjects that were injected with AAV-fDIO-Cre (orange points). For direct comparison, prior data from PV-Mecp2 WT (right black points) and PV-Mecp2 mutant (red points) are also provided. A one-way ANOVA of all groups revealed significant differences among them (F = 5.02, *p* < 0.01). Retrieval latencies were significantly longer in mice injected with fDIO-Cre as compared to control mice injected with AAV-GFP (n = 12 controls, latency: 0.230 ± 0.04; n = 12 mutants, latency: 0.381 ± 0.08; Sidak’s test, \**p* < 0.05).

We compared the pup retrieval performance on PND 0 of Flp-Flox subjects to that of GFP-Flox mice and found that mean retrieval latency was longer for Flp-Flox subjects (Figure 7B). A one-way ANOVA was used to compare PND 0 retrieval latencies between those groups and also between PV-Mecp2 mutant and PV-Mecp2 WT. Significant differences were detected among the groups (F = 5.02, *p* < 0.01) and post hoc testing detected significantly longer latencies for the Flp-Flox group (n = 12 Flp-Flox mice, n = 12 GFP-Flox mice; Sidak’s test, *p* < 0.05) and the PV-Mecp2 mutant group (n = 25 PV-Mecp2 mutant mice, n = 14 PV-Mecp2 WT mice; Sidak’s test, *p* < 0.001) when compared to their respective controls.

## DISCUSSION

Several lines of evidence from our previous work on *Mecp2^het^* mice strongly suggested that dysregulation of PVin in auditory cortex is a critical feature of the neuropathology underlying their failure to learn to perform pup retrieval behavior. Specifically, in the auditory cortices of *Mecp2^het^* virgin females co-housed with a dam and her litter, we observed dramatic overexpression of markers associated with PVin (parvalbumin protein and perineuronal nets) that are known to be antagonistic to plasticity (Pizzorusso et al., 2002; Carulli et al., 2010; de Vivo et al., 2013; Donato et al., 2013; Happel et al., 2014; Hou et al., 2017; Cisneros-Franco and de Villers-Sidani, 2019; reviewed in Rupert and Shea, 2022). This was accompanied by a lack of the disinhibitory plasticity found in the auditory cortex WT mice after exposure to pups (Lau et al., 2020). Several manipulations that ameliorated PV and PNN overexpression in *Mecp2^het^* subjects led to a resumption of behavior and partial restoration of the neural disinhibitory response (Krishnan et al., 2017; Lau et al., 2020). Here we present evidence that deletion of *Mecp2* selectively in PVin is sufficient to re-create many aspects of the neuropathology linked to the behavioral learning deficits that we observe in non-conditional, mosaic *Mecp2^het^* mutants.

### PVin have a disproportionate role in early establishment of retrieval behavior

Numerically speaking, cortical PVin make up a small population of neurons (accounting for about 10% of cortical neurons), yet they can powerfully affect neural activity (Cardin, 2018). Indeed, we compared the effects of deleting Mecp2 in PVin only with deleting it in two other major classes of cortical inhibitory neurons: somatostatin (SST) and vasoactive intestinal peptide (VIP) neurons. These populations are slightly less numerous than PVin but are of the same order of magnitude. We found no detectable effect on retrieval performance of selectively deleting *Mecp2* in SSTin or VIPin. This points to a specific function during retrieval for PV neurons, among all inhibitory subtypes, that makes the brain especially vulnerable to their loss of Mecp2.

Moreover, the admittedly short delay in the emergence of retrieval from PVin was no stronger or longer in mice that lacked Mecp2 in homeobox protein box (Emx1) neurons, which constitute ~88% of cortical neuron. This again suggests that the much smaller PVin population plays a specific and disproportionately large role in auditory cortical plasticity. Notably, in both PVin and Emx1 populations, removing *Mecp2* caused only a delayed emergence of retrieval behavior, not the sustained deficit we observed in non-conditional, mosaic *Mecp2^het^*. This suggests that *Mecp2* in PVin and Emx1 neurons each have an obligatory role in the early initiation of auditory cortical plasticity, but not necessarily in its subsequent maintenance. Yet, complete *Mecp2* deletion in either PVin or Emx1 neurons is less potent than mosaic absence of Mecp2 among all cell-type populations. This suggests that compensatory mechanisms involving nontargeted cell-types attenuate the effects of deleting *Mecp2* in only one cell type. Based on the results of deleting *Mecp2* in PVin during early adulthood, such compensatory mechanisms do not involve developmental processes. Moreover, suppression of typical expression patterns of PNNs acutely, just prior to introduction of pups, was sufficient to improve behavior within 5 d, despite any changes in the preceding developmental trajectory.

### Relationship of behavior to expression patterns and neurophysiology in PVin

PVin-specific deletion of *Mecp2* caused a very similar upregulation in PV and PNNs to that seen in mosaic *Mecp2^het^* mutants (Krishnan et al., 2017). However, unlike *Mecp2^het^* mice, the upregulation was a mix of transient and persistent increases. Specifically, PV-Mecp2 mutants exhibited a persistent increase in staining intensity of PV protein, yet the increase in staining intensity of PNNs present on PND 1 subsided by PND 5. It is possible that the failure to sustain high levels of PNN staining limits the duration of the disruption of behavior in PV-Mecp2 mutants. It is worth noting that because the changes in count of PVin and associated structures are so rapid (within 1 – 2 d), these changes very likely result from a change in expression intensity relative to our detection threshold, not a change in the absolute number of PVin cells themselves (i.e., celltype identity is unlikely to change over such a short time course).

Our prior work suggested that PNN expression is closely related to retrieval performance; not only were expression and performance correlated, but the administration of chondroitinase to dissolve PNNs in the auditory cortex actually improved behavior in *Mecp2^het^* (Krishnan et al., 2017). We therefore hypothesize that the long-term establishment of well-developed PNNs in the auditory cortex is a crucial barrier to the cortical plasticity underlying pup retrieval learning. Interestingly, deletion of *Mecp2* in PVin, while sufficient to establish more mature PNNs, is insufficient to sustain them. This implies that PNNs, despite preferentially surrounding PVin, are influenced by cell autonomous and non-cell autonomous processes on distinct timescales. In light of this, it will be interesting in future studies to see whether deletion of *Mecp2* in Emx1 neurons also lead to increased PNN expression at PV synapses, i.e. through a non-cell autonomous mechanism.

Our past work also revealed that maternal experience triggers disinhibited activity in the auditory cortex of WT females. In WTs this disinhibition was mediated by PVin but was abolished in PV-Mecp2 mutants (Lau et al., 2020). Here we find that in PV-Mecp2 mutants, PVin also do not decrease their stimulus-driven activity after the female acquires experience caring for pups. Since subjects in electrophysiology experiments were recorded after PND 6, and subjects in fiber photometry experiments were imaged through PND 5, the prevention of auditory cortical disinhibition outlasted the transient behavioral and molecular effects of Mecp2 deletion in PVin.

### Deletion of Mecp2 in PVin likely affects behavior by an acute and cell-autonomous mechanism

Importantly, our experiments with PV-specific Mecp2 knockout removed Mecp2 early in development and from all PV-expressing neurons throughout the brain. To bring greater spatiotemporal specificity to our manipulation, we adopted an intersectional strategy that required both Flp and Cre recombinases to remove Mecp2, allowing us to manipulate only PVin in the auditory cortex, and only in adulthood. We found that this also produced a transient delay in pup retrieval, establishing that the plasticity mechanisms that support that behavior also acutely require Mecp2 in adulthood, rather than during development alone. This observation argues in favor of the likelihood that the transient behavioral disruption is a cell-autonomous consequence of loss of Mecp2 in PVin of the auditory cortex because those are the same cells that are the effectors of the circuit disruption.

### Implications and future directions

A number of questions remain that should be the focus of future work. First, although our optical and electrical recordings from the auditory cortex were performed in awake animals, it is not yet known how the disinhibition we observe interacts with ongoing activity in freely behaving mice that are performing pup retrieval. PVin are important to the phenotype of *Mecp2^het^* mice, and they can modulate cortical activity from the single unit level up to more widespread features of brain state (Cardin, 2018), including gamma oscillations (Cardin et al., 2009; Sohal et al., 2009). Recording from actively retrieving mice may reveal unappreciated dynamic influences such as locomotor activity and arousal that may be mediated by PVin. Second, the relationship between PVin activity and construction of PNNs is not well understood. An interesting goal for future work will be to ascertain the relationship between PNNs and PVin activity, how they are affected by the activity of other cell-types, and on what timescale. Third, these questions about cell autonomous and non-cell autonomous influences of Mecp2 will be enlightened by targeted recordings from neurons that are individually identified as *Mecp^2+^* and *Mecp^2-^* in mosaic *Mecp2^het^* mice.

## Acknowledgments

The authors thank A. Zador, H. Hsieh, Z. J. Huang, and A. Banerjee for their thoughtful comments and guidance. The authors thank C. Nguyen, C. Kelahan, and R. Palaniswamy for technical support. The authors thank Rob Eifert and the CSHL machine shop staff for equipment support. This work was supported by a Royal Arch Masons Predoctoral Fellowship award from Autism Speaks to DDR and grants from the National Institutes of Mental Health (R01MH106656) and The Feil Foundation to SDS.

